# Cell Death Induced by Homoisoflavonoid Brazilin and its Semi-synthetic Derivates on MDA-MB-231 and MCF7 Breast Cancer Cell Lines

**DOI:** 10.64898/2026.01.09.698710

**Authors:** Miriam Zuñiga-Eulogio, Michael Quinteros, Alberto Hernández-Moreno, Tadeo Hernández-Moreno, Tapan Sharma, Mario Ordóñez, Carla Coste-Sanchez, Teresita Padilla-Benavides, Napoleón Navarro-Tito

**Affiliations:** Facultad de Ciencias Químico Biológicas, Universidad Autónoma de Guerrero, Chilpancingo, Gro, 39086, Mexico; Department of Molecular Biology and Biochemistry, Wesleyan University, Middletown, CT, 06459, USA; Centro de Investigaciones Químicas-IICBA, Universidad Autónoma del Estado de Morelos, Mor, 62209 Mexico; Department of Genetic and Cellular Medicine, University of Massachusetts Chan Medical School, Worcester, MA, 01605, USA; Department of Chemistry, Wesleyan University, Middletown, CT, 06459, USA

**Keywords:** Breast Cancer Treatment, Medicinal Chemistry, Flavonoids, Homoisoflavonoids, Brazilin, Semisynthesis, Cell death

## Abstract

Flavonoids are naturally occurring polyphenolic compounds that have been extensively explored as scaffolds for drug development due to their diverse biological activities. Brazilin, a homoisoflavonoid with reported antitumoral properties, does not fully meet pharmaceutical criteria, and chemical modification of natural compounds is often required to enhance bioactivity and efficacy. Here, we evaluated the pro-apoptotic activity of Brazilin and its semi-synthesized methoxylated (OMe)_3_ and acetylated (OAc)_3_ derivatives in triple-negative MDA-MB-231 and luminal A MCF7 breast cancer cell lines.

We assessed cell viability, proliferation, oxidative stress, and mitochondrial integrity, and analyzed apoptotic features using confocal microscopy, western blotting, and RT-qPCR. In addition, RNA sequencing was performed to characterize transcriptomic changes in MDA-MB-231 cells following treatment with unmodified Brazilin or its derivatives. Brazilin and Brazilin-(OAc)_3_ significantly reduced cell viability and proliferation in MDA-MB-231 cells, whereas MCF7 cells exhibited increased viability and growth in response to Brazilin-(OMe)_3_. In MDA-MB-231 cells, treatment with Brazilin and Brazilin-(OAc)_3_ induced apoptosis-associated features, including chromatin condensation, γH2AX accumulation, and PARP cleavage. These effects were accompanied by a modest increase in mitochondrial oxidative stress and loss of mitochondrial membrane potential. Notably, no cytotoxic or apoptotic features were detected in non-tumorigenic MCF10A cells.

Transcriptomic analysis revealed that Brazilin treatment upregulated genes associated with endoplasmic reticulum stress, including *ATF3*, in MDA-MB-231 cells. Collectively, our results indicate that Brazilin and its acetylated derivative selectively induce mitochondrial stress and cell death in triple-negative breast cancer cells, potentially involving ER stress pathways.

## INTRODUCTION

In cancer therapy, around 57% of currently used chemotherapeutics have been inspired or derived from natural compounds. Therefore, we continue looking for new leads for drug development in secondary metabolites of fruits and plants^1, 2^. Among the natural compounds, flavonoids have emerged as a diverse group of bioactive metabolites. Flavonoids are diet-abundant polyphenolic compounds, classified according to their complexity and number of saturations to the core structure. Among these, homoisoflavonoids (3-benzyl-chroman-4-ones) are a subclass of flavonoids^3^, subclassified into scillascillin-, caesalpin-, sappanin-, protosappanin and Brazilin-type^4, 5^. These compounds are well-known for their anti-inflammatory, anti-microbial, anti-diabetic, antioxidant and anti-tumoral bioactivities^4, 6, 7^. The anti-tumoral activity of several homoisoflavonoids has been studied in models of lung cancer^8–10^, ovarian cancer, melanoma, non-small cell lung cancer^11^, colorectal cancer, leukemia^12^, liver cancer and breast cancer^8, 13–15^.

Brazilin ((6*aS*,11*bR*)-7,11*b*-dihydro-6*H*-indeno[2,1-c]chromene-3,6*a*,9,10-tetrol; CAS 474-07-7) is an homoisoflavonoid isolated from *Caesalpinia sappan* and *Haematoxylum brasiletto* from Southeast Asia and Pacific coast area, respectively^5, 16, 17^. In traditional ethnopharmacology, numerous plant parts of *C. sappan* and *H. brasiletto* species are frequently used in infusions to address health issues such as fever, inflammation, dysentery, diabetes, pneumonia, anemia, and hypertension^18^. Additionally, Brazilin extracts are blended into balms to alleviate swelling, rheumatism, and wounds, among other therapeutic uses^18^. Furthermore, purified Brazilin inhibits proliferation and cell migration and promotes apoptosis of *in vitro* models of colon cancer, lung cancer, bladder cancer, cervical cancer, glioblastoma, and osteosarcoma^19–24^.

Apoptosis evasion from tumor cells is related to initiation, cancer progression, and chemoresistance^25, 26^. Apoptosis is driven by the death receptor-dependent extrinsic or the mitochondrial-dependent intrinsic pathways; however, endoplasmic reticulum (ER) stress can also lead to cell death^27^. The receptor extrinsic pathway begins with the binding of ligands (TRAIL, TNF, FasL) to the respective death receptors (DR4/5, TNFR, and Fas)^28^. Intracellular adaptor FADD recruits pro-caspase 8/10 to form the death-inducing signaling complex (DISC), which activates the effector caspases 3/7^28^. In the mitochondrial intrinsic pathway, the anti-apoptotic Bcl-2 and Bcl-X_L_ are outnumbered by pro-apoptotic Bax and Bak proteins, which promote mitochondrial outer membrane permeability (MOMP) to release cytochrome C (Cyt C); Cyt C binds Apaf-1 and pro-caspase 9 to form the apoptosome, then the effector caspases 3/7 are activated^28^. Apoptosis is a finely regulated process that involves the participation of phagocytic cells, which guarantees the integrity of adjacent cells and tissues. Thus, understanding the subjacent mechanisms that drive cellular death resistance in cancer cells and developing specific and effective therapy against these remarkable targets^29, 30^.

In cancer therapy, the approaches targeting apoptosis have been mainly focused on inhibiting the anti-apoptotic members of the Bcl-2 (B-Cell Lymphoma 2) family and IAPs (inhibitor of apoptosis proteins); mimetics of SMAC (second mitochondrial activator of caspases) proteins; use of death receptor agonists; and enhancement of p53 activity by inhibiting MDM2 (Mouse Double Minute 2)^31^. A less common strategy to induce apoptosis consists of promoting the activation of the ATF4-CHOP-DR5 axis, which further enhances the extrinsic apoptosis pathway^31–33^. Moreover, ATF4 (Activating Transcription Factor 4) activation is also involved in endoplasmic reticulum stress (ER stress) response-mediated apoptosis; the unfolded protein response (UPR) sensor PERK, located in the ER membrane, activates eIF2α, which promotes the translation of ATF4. ATF4 induces the expression of anti-oxidative genes and pro-apoptotic genes such as *BIM* and *DR5*^34^. ER stress-dependent apoptosis also involves the activation of alternative proteases like calpains and caspase 4, activating the JNK stress-response pathway and the Ca^2+^ release-induced loss of mitochondrial integrity^35^.

Our group recently isolated and purified the homoisoflavonoid Brazilin from the heartwood of *H. brasiletto* and then synthesized two semi-synthetic derivatives by the addition of three methyl or acetyl groups to 3’, 9’, and 10’ OH groups of Brazilin: Brazilin-(OMe)_3_ and Brazilin-(OAc)_3_, respectively^36^. Here, we assessed their pro-apoptotic activity in the triple-negative breast cancer (TNBC) MDA-MB-231, luminal A (LA) breast cancer MCF7 cells, and the non-tumorigenic breast epithelial MCF10A cell line. We analyzed cell viability using (4,5-dimethylthiazol-2-yl)-2,5-diphenyltetrazolium bromide (MTT) and Trypan blue exclusion assays. Confocal microscopy, western blot, and RT-qPCR analyses evaluated apoptotic features, and DHE, MitoSox, and TMRE standard assays assessed oxidative stress and mitochondrial integrity. RNA-seq analyses were performed in MDA-MB-231 cells to understand transcriptional changes derived from Brazilin and derivatives treatments. Our study demonstrated that both Brazilin and Brazilin-(OAc)_3_ significantly affected the cell viability and growth of MDA-MB-231 cells. The results revealed DNA damage, activation of caspase 3, and PARP cleavage, irrespective of the expression levels of Bax/Bak and Bcl-2/Bcl-X_L_. In contrast, MCF7 and non-tumorigenic MCF10A cells did not exhibit a cytotoxic response to Brazilin or derivatives. RNA-seq analysis indicated the involvement of the ER stress-response *ATF3* gene, and the loss of mitochondrial integrity appeared to contribute to the apoptosis mechanism induced by Brazilin. Our findings suggest that the death observed in MDA-MB-231 cells following treatment with Brazilin and Brazilin-(OAc)_3_ is closely linked to the ER stress response.

## RESULTS AND DISCUSSION

### Brazilin and its acetylated derivative selectively reduce viability and proliferation of TNBC cells

Cell viability is a primary parameter for assessing the antitumoral potential of natural products, reflecting either cytostatic or cytotoxic effects. Homoisoflavonoids, including Brazilin, have previously demonstrated selective cytotoxicity across cancer cell lines, although some others exhibit phytoestrogenic effects in hormone-responsive cells. To investigate the antitumor activity of Brazilin and two semi-synthetic derivatives, Brazilin-(OMe)_3_ and Brazilin-(OAc) _3,_ we evaluated their effects on triple-negative MDA-MB-231, luminal A MCF7, and non-tumorigenic MCF10A cells. Treatment with 20 µM Brazilin or Brazilin-(OAc)_3_ for 48 h significantly reduced MDA-MB-231 viability (IC_50_ = 40.73 ± 1.10 µM and 37.26 ± 1.32 µM, respectively), whereas Brazilin-(OMe)_3_ only affected viability at ≥80 µM (**Figure 1a**; **Table 1**). MCF7 cells exhibited reduced viability at ≥40 µM for Brazilin and Brazilin-(OAc)_3_, with the methoxylated derivative slightly increasing proliferation at low concentrations (**Figure 1b**). Non-tumorigenic MCF10A cells were less sensitive, with IC_50_ values of 46.09 ± 1.17 µM (Brazilin) and 55.93 ± 1.66 µM (Brazilin-(OAc)_3_), whereas Brazilin-(OMe)_3_ again showed minimal effects (**Figure 1c**, **Table 1**). These data indicate selective cytotoxicity toward TNBC cells, a feature that is clinically advantageous to minimize off-target toxicity.

**Figure 1.**
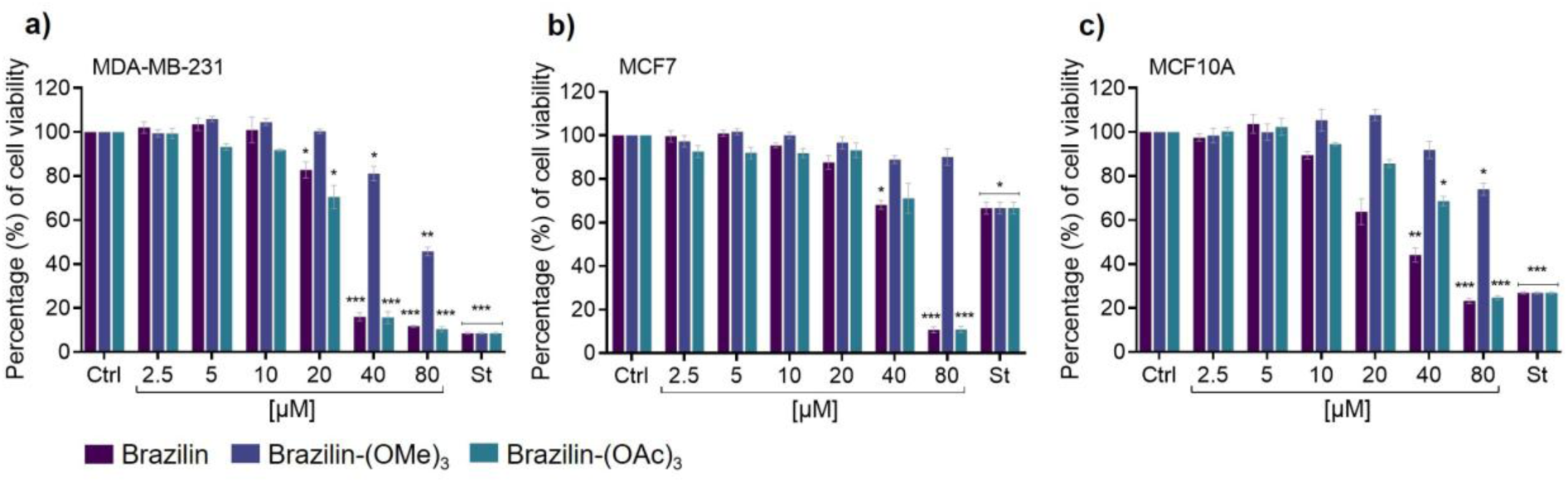
Effect of brazilin and derivatives on cell viability of breast cancer and non-tumorigenic epithelial cells. MDA-MB-231, MCF7, and MCF10A cells were treated for 48 h with 2.5-80 µM of brazilin, brazilin-(OMe)_3_, or brazilin-(OAc)_3_. Cell viability was assessed using an MTT assay (0.5 mg/mL). Staurosporine (St, 50 nM) served as a positive control. Viability is expressed as a percentage relative to untreated control cells (Ctrl) for **(a)** MDA-MB-231, **(b)** MCF7, and **(c)** MCF10A. Data represent mean ± SD from three independent biological replicates. Statistical significance: *p < 0.05, **p < 0.01, ***p < 0.001.

**Table 1.** IC_50_ of brazilin and derivatives in breast cancer and epithelial non-tumorigenic cells.

|  | ***Brazilin*** | ***Br-(OMe)<sub>3</sub>*** | ***Br-(OAc)<sub>3</sub>*** |
| --- | --- | --- | --- |
| *MDA-MB-231* | 40.73±1.10 µM | 79.31±5.03 µM | 37.26±1.32 µM |
| *MCF7* | 49±2.18 µM | >80 µM* | 49.97±3.91 µM |
| *MCF10A* | 46.09±1.17 µM | >80 µM* | 55.93±1.66 µM |
| *IC<sub>50</sub> out of the concentration range for estimation |  |  |  |

Long-term proliferation assays further confirmed selectivity (**Figure 2**). At 96 h post-treatment with 20 µM compounds, MDA-MB-231 cell numbers were reduced by 76% and 82% with Brazilin and Brazilin-(OAc)_3_, respectively, while Brazilin-(OMe)_3_ caused only a 28% reduction (**Figure 2a, d**). In contrast, proliferation of MCF7 and MCF10A cells was only modestly affected (**Figure 2b-d**). Morphological analysis revealed rounding and reduced density in MDA-MB-231 cells treated with active compounds, whereas MCF7 and MCF10A cells retained largely normal morphology (**Figure 2d**). At sub-toxic concentrations (2.5-5 µM), Brazilin and Brazilin-(OAc)_3_ increased proliferation in MDA-MB-231 cells, consistent with hormetic dose responses observed for polyphenols, whereas the methoxylated derivative had minimal effect. As expected, 40 µM treatments completely prevented the proliferation of MDA-MB-231 cells (**Supp. Figure 1c-d**). These results highlight the importance of OH-substitution patterns for bioactivity, with acetylation preserving or enhancing cytotoxicity while methylation abolished it. Notably, low-concentration proliferative effects of Brazilin-(OMe)_3_ in MCF7 cells (**Supp. Figure 1f-g)** may reflect phytoestrogenic signaling mediated by the estrogen receptor α (ERα), consistent with prior reports of differential flavonoid activity across hormone-responsive lines.

**Figure 2.**
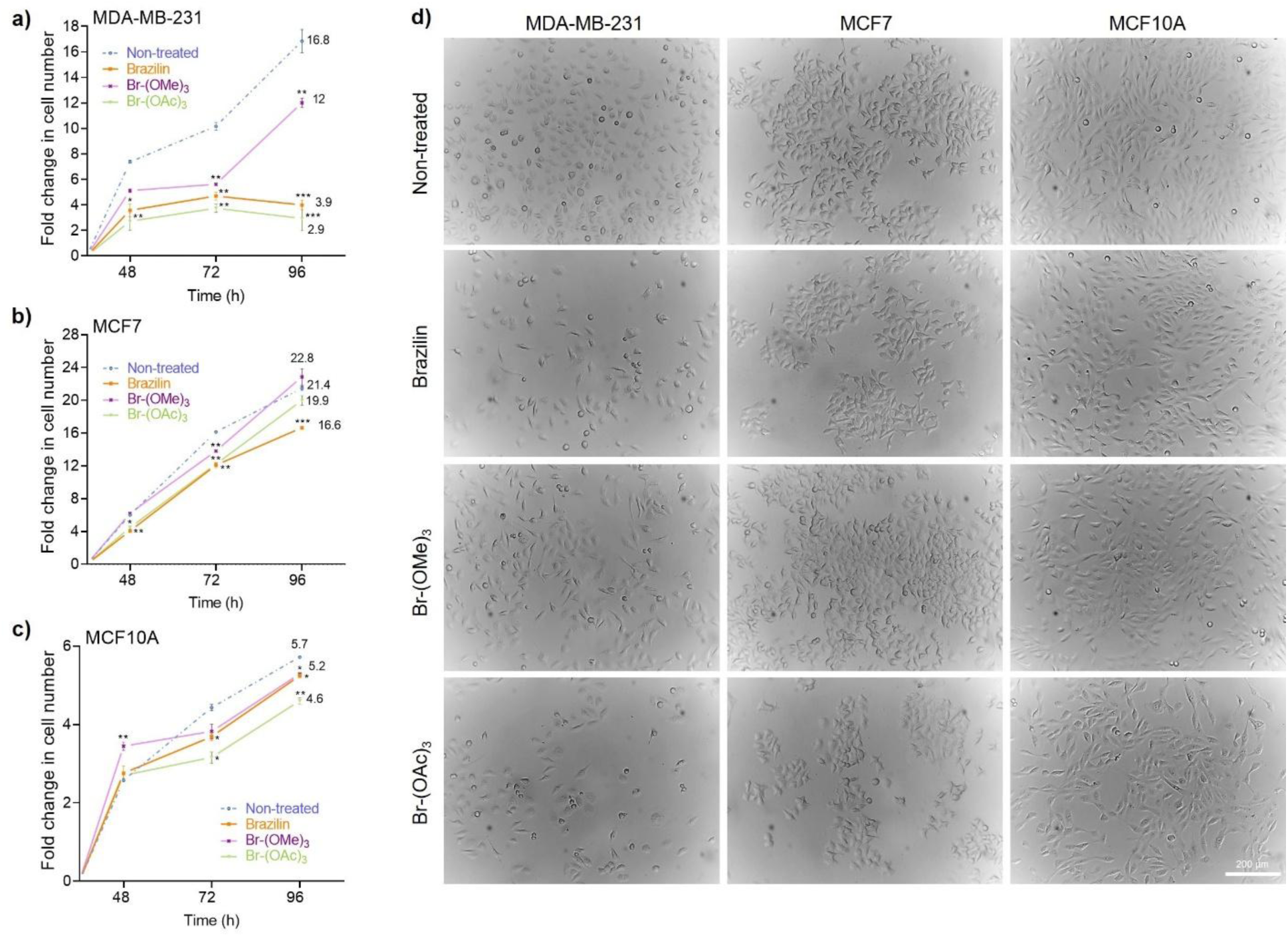
Brazilin and derivatives alter proliferation and morphology of breast cancer and non-tumorigenic epithelial cells. Proliferation of **(a)** MDA-MB-231, **(b)** MCF7, and **(c)** MCF10A cells treated with 20 µM of brazilin or derivatives was assessed by trypan blue cell counting at 48, 72, and 96 h. Data are expressed as fold change in cell number over time and shown as mean ± SE; mean values at 96 h are indicated. See **Supp** Figure 1c-k for complete cell growth data at 2.5, 5, and 40 µM. Statistical significance: *p < 0.05, **p < 0.01, ***p < 0.001. **(d)** Representative brightfield images of cells after 96 h of treatment (20 µM) captured using a NIKON® ECLIPSE Ts2 microscope. Scale bar = 200 µm. See **Supp** Figure 2 for images at 48 and 72 h. Data represent three independent biological replicates.

### Brazilin and its derivatives induce apoptotic features in MDA-MB-231 cells

To investigate whether reduced viability resulted from apoptosis, MDA-MB-231, MCF7, and MCF10A cells were treated with 20 µM of each compounds for 48 h. Caspase-3 activity and PARP cleavage were selectively induced in MDA-MB-231 cells by Brazilin and Brazilin-(OMe)_3_, whereas MCF7 cells showed only modest activation and MCF10A cells remained largely unaffected (**Figure 3a-c**). Confocal microscopy revealed strong nuclear condensation and γH2AX foci in MDA-MB-231 cells treated with Brazilin, indicative of DNA damage and apoptotic execution (**Figure 4a**). Brazilin-(OAc)_3_ induced γH2AX foci without pronounced nuclear condensation, suggesting cell death may be mediated through alternative stress pathways (**Figure 4a**). Analysis of Bcl-2 family expression revealed modest increases in Bax protein with Brazilin-(OMe)_3_ and slight Bcl-2 elevation with Brazilin-(OAc)_3_ in MDA-MB-231 cells (**Supp. Figure 3e-g**). MCF7 and MCF10A cells exhibited limited changes. These findings indicate that apoptosis induction is not strictly dependent on canonical Bax/Bak activation and may involve mitochondrial and ER stress-mediated pathways.

**Figure 3.**
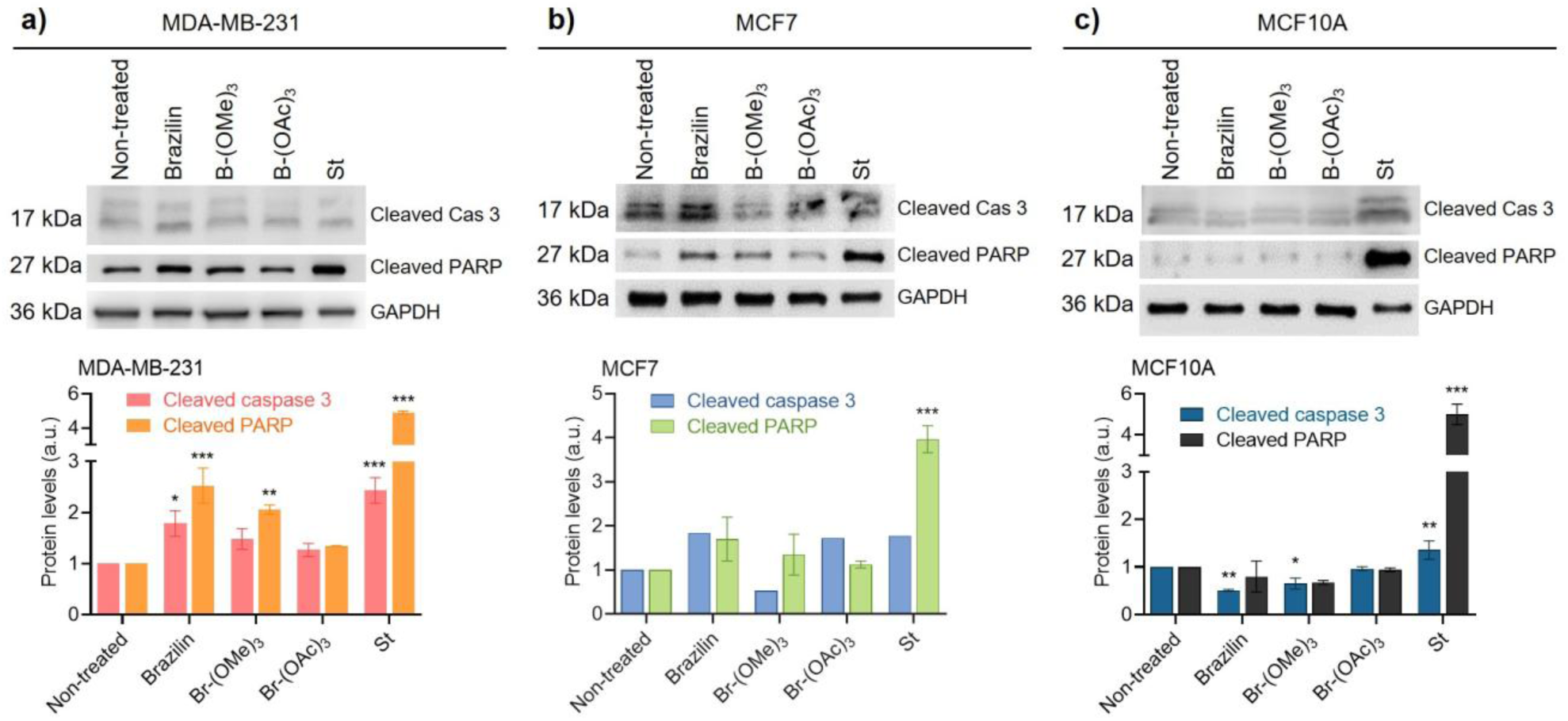
Brazilin and derivatives induce caspase 3 activation and PARP cleavage in MDA-MB-231 cells. Representative immunoblots (top) and quantification (bottom) of active cleaved caspase 3 and cleaved PARP in **(a)** MDA-MB-231, **(b)** MCF7, and **(c)** MCF10A cells treated with 20 µM of brazilin or derivatives for 48 h. Staurosporine (50 nM) was used as a positive control for apoptosis. GAPDH was used as a loading control. Data are presented as mean ± SE from independent biological replicates. Statistical significance: *p < 0.05, **p < 0.01, ***p < 0.001.

**Figure 4.**
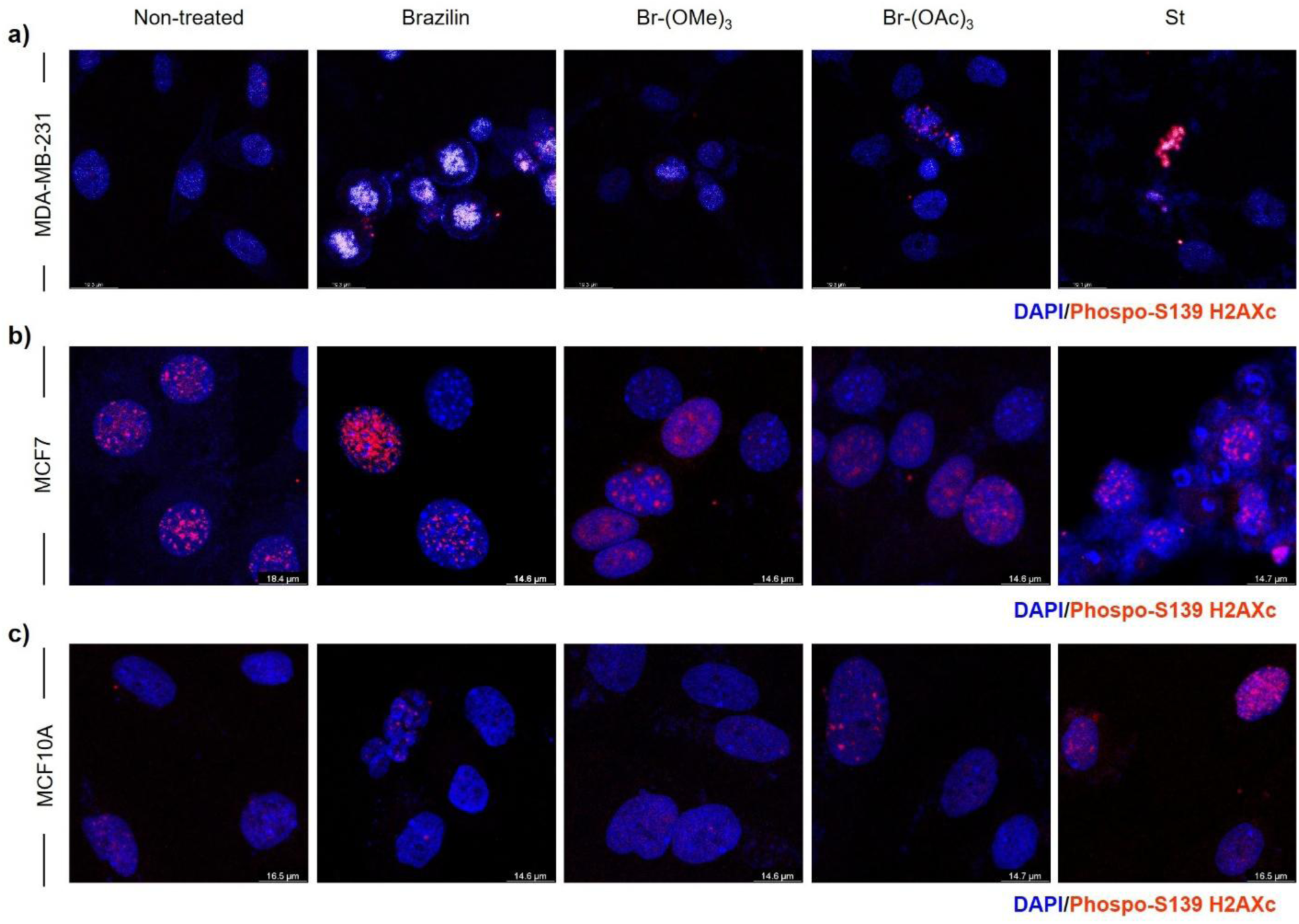
Brazilin and derivatives induce DNA damage and chromatin condensation in breast cancer cells. Confocal images of **(a)** MDA-MB-231, **(b)** MCF7, and **(c)** MCF10A cells treated with 20 µM of brazilin, brazilin-(OMe)_3_, or brazilin-(OAc)_3_ for 48 h. DNA damage is indicated by γH2AX (phospo-S139 H2AXc) immunostaining (red), and chromatin condensation is visualized by intense DAPI staining (blue). Scale bar = 14-19 µm (see individual images). Images were acquired using a Leica SP8 confocal microscope with Leica Application Suite X. Representative images from at least three independent experiments.

### Selective mitochondrial stress and transcriptional responses underlie TNBC cytotoxicity

Homoisoflavonoids can act as antioxidants or pro-oxidants depending on cellular context. Brazilin exhibited robust radical-scavenging activity in the DPPH assay (20 µM scavenging 29.7 ± 5.4%), whereas derivatives displayed minimal antioxidant capacity. We estimated the half-maximal effective concentration (EC_50_) to scavenge DPPH radicals at >500 µM for brazilin-(OMe)_3_, and >1 mM for brazilin-(OAc)_3_ (**Figure 5a**; **Table 2**). Despite this, both Brazilin and Brazilin-(OAc)_3_ induced mild cytoplasmic ROS in MDA-MB-231 cells (**Figure 5b**) and selectively induced mitochondrial oxidative stress (**Figure 5c**) and compromised mitochondrial membrane potential at higher concentrations (40 µM), while non-tumorigenic cells remained largely unaffected (**Figure 5c-h**). These results suggest selective mitochondrial vulnerability in TNBC cells, potentially amplified by elevated basal ROS.

**Figure 5.**
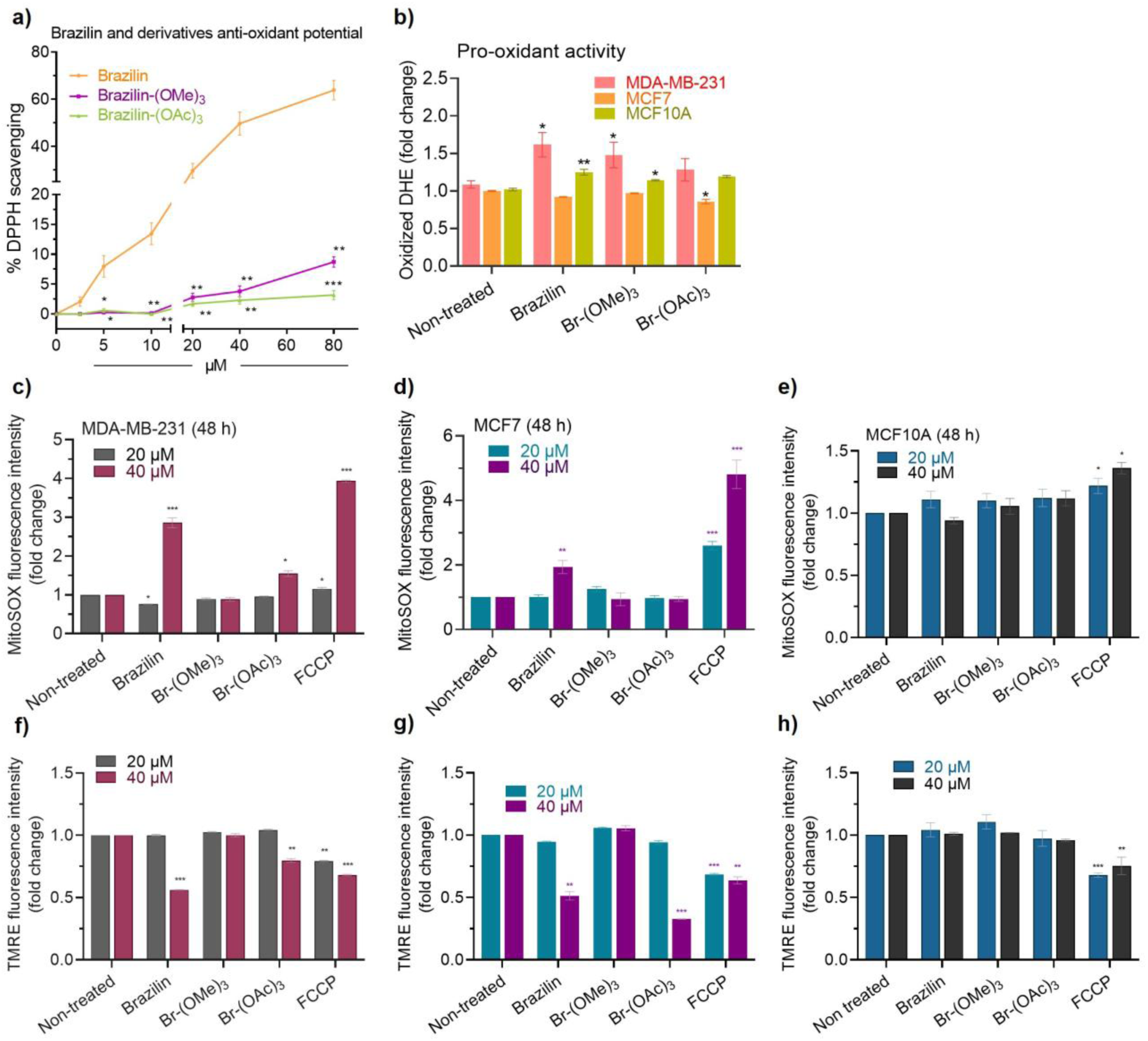
Antioxidant potential and pro-oxidant effects of brazilin and derivatives in breast cancer and non-tumorigenic cells. **(a)** Radical scavenging activity of brazilin, brazilin-(OMe)_3_, and brazilin-(OAc)_3_ (2.5-80 µM) was assessed using DPPH synthetic radicals. **(b)** Cytoplasmic ROS levels were measured in MDA-MB-231, MCF7, and MCF10A cells after 48 h treatment with 20 µM of each compound by fluorescent detection of DHE oxidation. **(c-h)** Mitochondrial ROS (MitoSOX) and mitochondrial membrane potential (TMRE) were evaluated in (c,f) MDA-MB-231, **(d,g)** MCF7, and **(e,h)** MCF10A cells treated for 48 h with 20 µM or 40 µM of brazilin or derivatives. FCCP was used as a positive control for mitochondrial depolarization. Fluorescence intensities are plotted as mean ± SE relative to untreated cells for each concentration. *p<0.05, **p<0.001, ***p<0.001. Data represents three independent biological replicates.

**Figure 6.**
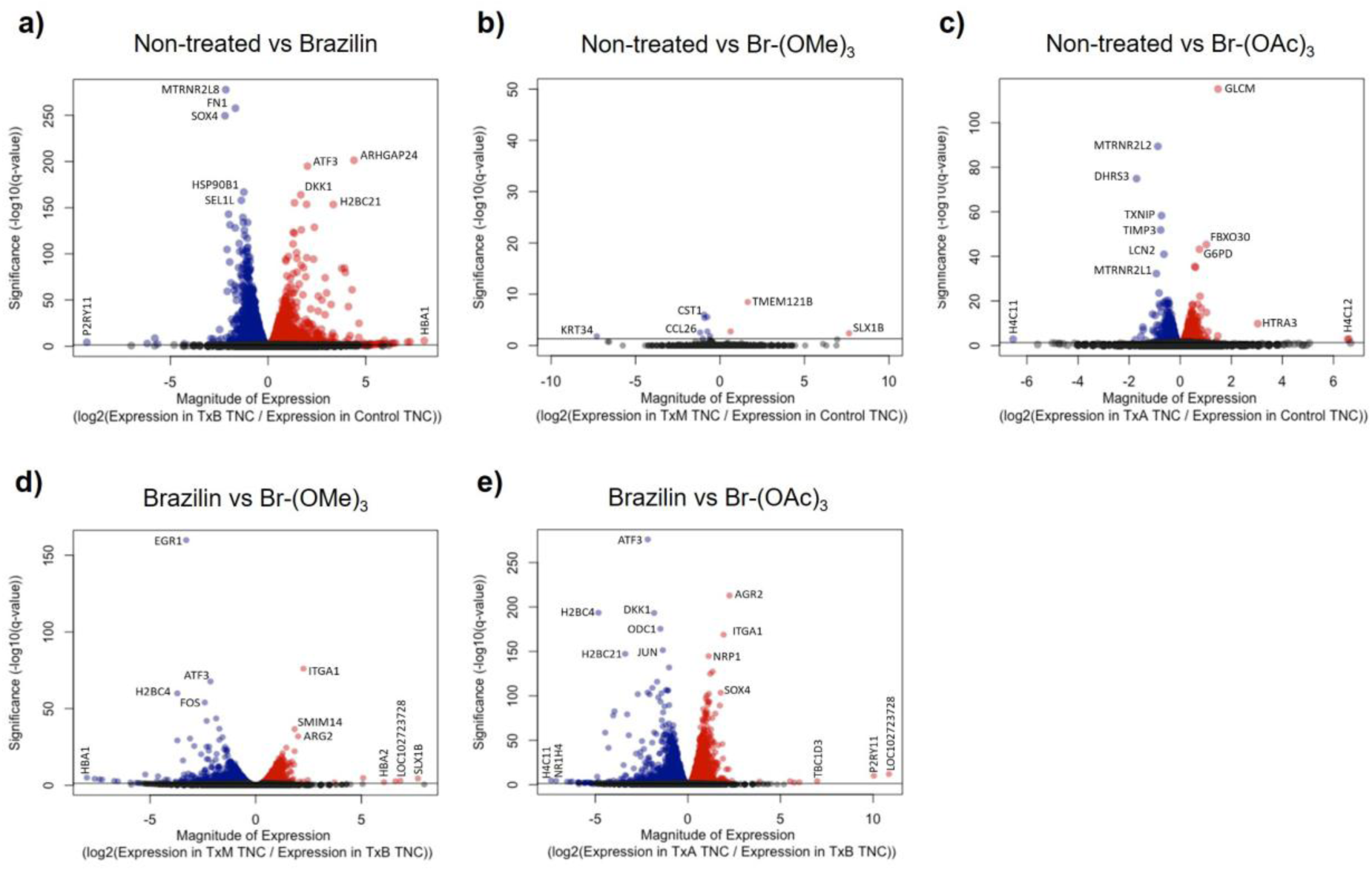
Transcriptomic changes in MDA-MB-231 cells treated with brazilin and derivatives. Volcano plots showing differentially expressed genes between untreated cells and cells treated with **(a)** brazilin, **(b)** brazilin-(OMe)_3_, or **(c)** brazilin-(OAc)_3_. Comparisons between brazilin and its derivatives are shown in **(d)** brazilin vs. brazilin-(OMe)_3_ and **(e)** brazilin vs. brazilin-(OAc)_3_. The y-axis represents -log_10_(p-value), and the x-axis shows log_2_ fold change. Red and blue dots indicate significantly up- and downregulated transcripts (FDR < 0.05), respectively, while gray dots represent transcripts that did not reach statistical significance (FDR > 0.05).

**Table 2.** EC_50_ of radical scavenger capacity of brazilin and derivatives.

| **Brazilin** | **Brazilin-(OMe)<sub>3</sub>** | **Brazilin-(OAc)<sub>3</sub>** |
| --- | --- | --- |
| 47.2±1.76 µM | 539.48±10.15 µM | >1 mM |

RNA-seq analysis revealed transcriptomic perturbations consistent with stress-induced apoptosis. Brazilin modulated 2,885 genes, downregulating cilium organization and ER stress-response genes and upregulating rRNA/ncRNA processing and extracellular matrix (ECM) organization genes (**Figure 7a**). Brazilin-(OAc)_3_ altered 193 genes, promoting apoptosis-regulatory gene expression (e.g., *GCLM*) and downregulating anti-apoptotic humanin-encoding genes (*MTRNR2L 1/2/8*; **Figure 7b**; **Figure 8e-g**). The methoxylated derivative induced minimal transcriptional changes, consistent with its weak phenotypic effects. RT-qPCR validation confirmed upregulation of stress response genes (*ATF3, H2BC21*) and downregulation (*SOX4, MTRNR2L 1/2*) of pro-survival genes in MDA-MB-231 cells (**Figure 8a-d**).

**Figure 7.**
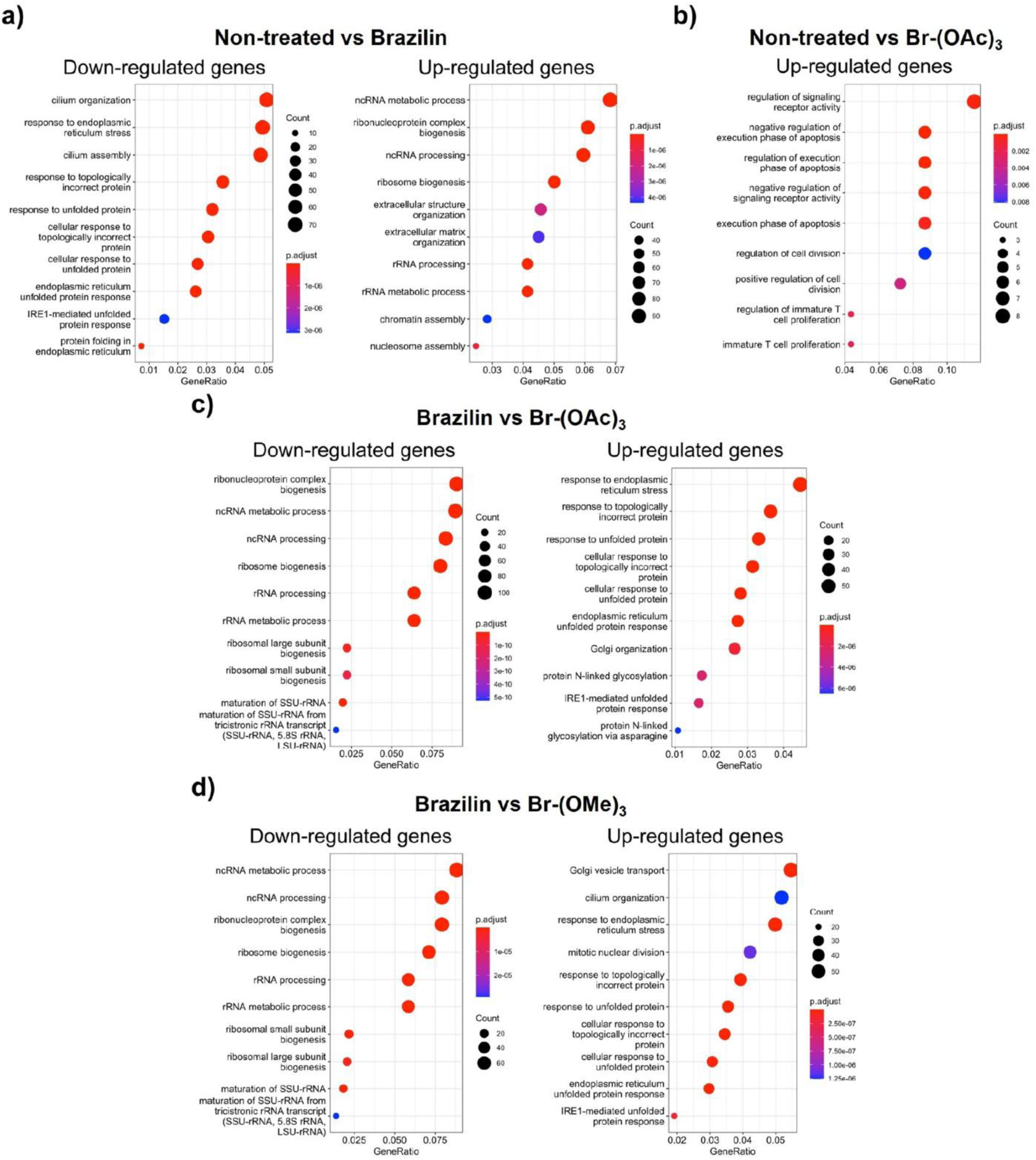
Gene ontology (GO) analysis of differentially expressed genes in MDA-MB-231 cells treated with brazilin and derivatives. GO term enrichment analysis comparing **(a)** untreated control versus brazilin-treated cells, **(b)** control versus brazilin-(OAc)_3_-treated cells, **(c)** brazilin versus brazilin-(OAc)_3_, and **(d)** brazilin versus brazilin-(OMe)_3_. The cut-off for significance was set at -log_10_(adjusted p-value) ≥ 2.0. See **Supp. Table S2** for the full list of differentially expressed genes and associated GO terms.

**Figure 8.**
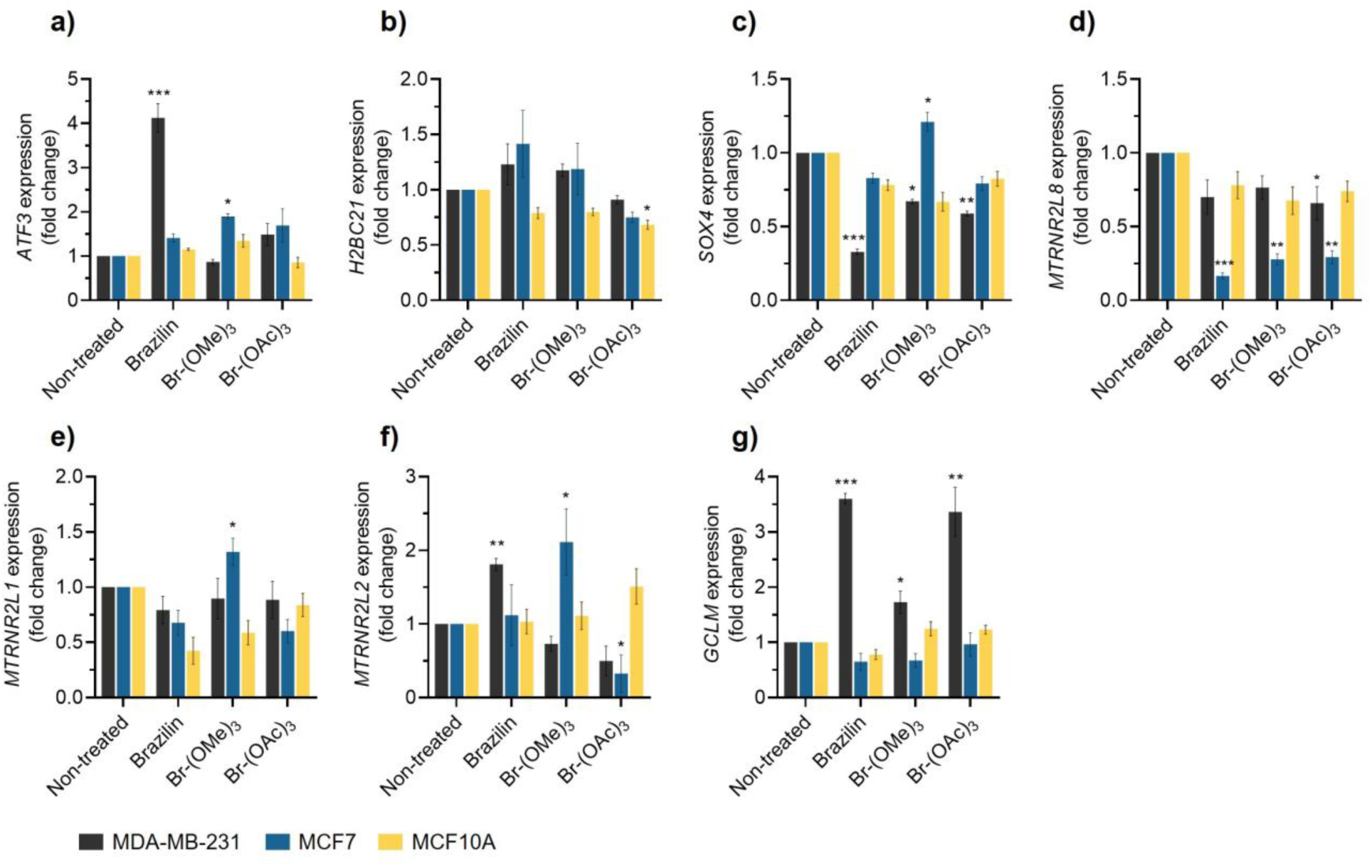
Validation of differentially expressed genes in breast cancer and non-tumorigenic epithelial cells treated with brazilin and derivatives. Differentially expressed genes (DEG) identified by RNA-seq in MDA-MB-231 cells. (a) *ATF3*, (b) *H2BC21*, (c) *SOX4*, (d) *MTRNR2L8*, (e) *MTRNR2L1*, (f) *MTRNR2L2*, and (g) *GCLM*, were validated by RT-qPCR in MDA-MB-231, MCF7, and MCF10A cells treated with 20 µM brazilin or derivatives for 48 h. Data are expressed as mean ± SE from three independent biological replicates. *p < 0.05, **p < 0.001, ***p < 0.001.

Collectively, these data support a model in which Brazilin and its acetylated derivative induce TNBC cell death through a combination of mild oxidative stress, selective mitochondrial dysfunction, and ER stress-linked transcriptional reprogramming that converges on cell death. Methylation abolishes these effects, underscoring the critical role of OH-substitution for bioactivity. Our study identifies Brazilin as a selective anti-tumoral homoisoflavonoid against TNBC cells, with the acetylated derivative enhancing cytotoxic efficacy while sparing non-tumorigenic cells. Mechanistically, cytotoxicity involves selective mitochondrial stress, induction of stress-response genes, and activation of apoptosis, with limited engagement of canonical Bax/Bak pathways. These findings highlight acetylation as a promising chemical diversification strategy to optimize flavonoid bioactivity and selectivity, warranting further preclinical evaluation.

## EXPERIMENTAL SECTION

### Reagents and antibodies

3-[4,5-dimethylthiazol-2-yl]-2,5 diphenyl tetrazolium bromide (MTT; M2128), 2,2-diphenyl-1-picrylhydrazyl (DPPH; D9132) and dihydroethidium (DHE; 37291) were purchased from Sigma-Aldrich (St Louis, MO). MitoSOX^TM^ (M36008) red mitochondrial superoxide indicator was obtained from Invitrogen-Thermo Fisher Scientific (Waltham, MA), the tetramethyl-rhodamine methyl ester (TMRE)-Mitochondrial Membrane Potential Assay Kit (ab113852) from Abcam (Cambridge, MA). Primary antibodies anti-Bax (A0207), anti-Bcl-2 (A19693), anti-cleaved-PARP (A19612), anti-GAPDH (A19056), and anti-phospho-Histone H2AX S139 (γH2AX; AP0099) were purchased from ABclonal (Woburn, MA), and for anti-cleaved-caspase 3 (D175) (#9664) from Cell Signaling (Danvers, MA). Secondary antibody anti-rabbit HRP (31-460) from Thermo Fisher Scientific. The secondary antibody Alexa Fluor 633 goat anti-rabbit (A21070) was from Invitrogen-Thermo Fisher Scientific (Waltham, MA) and Vectashield Plus Antifade Mounting Medium with DAPI (H-2000) from VectorLaboratories (Newark, CA).

### Brazilin and semi-synthetic derivatives

Brazilin ((6a*S*,11b*R*) -7,11b-dihydro-6*H*-indeno[2,1-*c*]chromene-3,6a,9,10-tetrol; CAS 474-07-7) isolation and purification from the duramen of *H. brasiletto*, as well as the semi-synthesis workflow to obtain the derivatives Brazilin-(OMe)_3_ ((6a*S*,11b*R*)-3,9,10-trimethoxy-7,11b-dihydroindeno[2,1-c]chromen-6a(6*H*)-ol) and Brazilin-(OAc)_3_ ((6a*S*,11b*R*)-6a-hydroxy-6,6a,7,11b-tetrahydroindeno[2,1-*e*]chromene-3,9,10-triyl triacetate) were previously described^36^. Dr. Jorge Bello-Martínez (Universidad Autónoma de Guerrero) provided us with the initial aliquot of Brazilin isolated and purified from *H. brasiletto*. Brazilin-(OMe)_3_ and Brazilin-(OAc)_3_ were synthesized in the Design and Synthesis of Bioactive Compounds Laboratory, under the direction of Dr. Mario Ordóñez (Universidad Autónoma del Estado de Morelos). The chemical structures of Brazilin, Brazilin-(OMe)_3_, and Brazilin-(OAc)_3_, as described before^36^, are shown in **Supp Figure 4**.

### Cell culture and treatments

Mammary gland-derived cell lines MCF10A (CRL-10317™), MDA-MB-231 (CRM-HTB-26™), and MCF7 (HTB-22™) were purchased from the American Type Culture Collection (ATCC, Manassas, VA). Breast cancer MCF7 and MDA-MB-231 cells were cultured in Culture Dulbecco’s Modified Eagle Medium (DMEM; Sigma-Aldrich) with 5% fetal bovine serum (FBS) and 1% penicillin G/streptomycin (GIBCO). The non-tumorigenic mammary epithelial MCF10A cell line was cultured in DMEM supplemented with 20 ng/mL recombinant EGF 0.5 µg/mL hydrocortisone, 10 µg/mL insulin, 10% FBS, and 1% penicillin/streptomycin. All cultures were maintained in a humified atmosphere containing 5% CO_2_ at 37°C. For experimental purposes, cells were rinsed with sterile PBS before any treatment. Different concentrations (2.5 µM, 5 µM, 10 µM, 20 µM, 40 µM, and 80 µM) of Brazilin and derivatives Brazilin-(OMe)_3_ and Brazilin-(OAc)_3_ were administrated for 48 h, 72 h, or 96 h. 50 nM Staurosporine was used as positive control of apoptosis, and DMSO was used as a treatment-solvent control. Treatment was finished by removing the media.

### Cell viability assay

Cell viability was measured by 3-[4,5-dimethylthiazol-2-yl]-2,5 diphenyl tetrazolium bromide (MTT) assay^37, 38^. MDA-MB-231, MCF7, or MCF10A cells were harvested and counted, 2x10^4^ cells were seeded in a 96-well microtiter plate, then treatments with Brazilin and derivatives were performed as described above. After 24 and 48 h of treatment, the media was removed, then 0.5 mg/mL MTT was added to each well and incubated protected from light at 37°C for 4 h. Formazan crystals were dissolved in DMSO and absorbances were measured at 570 nm in a Multiskan^TM^ GO microplate spectrophotometer (Thermo Fisher Scientific). The percentage of cell viability was calculated as (absorbance of samples/absorbance of control) x100, then the half maximal inhibitory concentration (IC_50_) was calculated by non-linear regression. IC_50_ was used to compare all compounds’ cytotoxic effects and determine the drug sensitivity in our cultured models.

### Cell Proliferation Assays

MDA-MB-231, MCF7, and MCF10A 1x10^4^ cells/cm^2^ were seeded in 24 well plates with different concentrations (2.5-40 µM) of Brazilin and its derivatives. Cells were collected at 48, 72, and 96 h after seeding with treatments. Cells were trypsinized and stained with 0.4% trypan blue, and the number of live cells was measured using a Spectrum Cellometer (Nexcelom Biosciences, Lawrence, MA). Data was processed with FCS Express 7 software (De Novo Software, Dotmatics).

### Brightfield and confocal microscopy

At each point of the proliferation assays, brightfield images of MDA-MB-231, MCF7, and MCF10A cells were obtained using a NIKON ® ECLIPSE Ts2 microscope. For confocal microscopy, MDA-MB-231, MCF7, and MCF10A cells were seeded in coverslips and treated with 20 µM Brazilin and derivatives for 48 h, then fixed with 10% formalin-PBS and permeabilized with 0.5% Triton X-100. The primary antibody phospo-H2AX S139 (γH2AX) (1:250 dilution) was incubated overnight in 3% FBS-PBS solution, and the secondary antibody Alexa Fluor 633 anti-rabbit (1:250 dilution) was incubated for 2 h protected from light. Coverslips were mounted using VECTASHIELD with DAPI, imaged with a Leica SP8 Confocal Microscope, and analyzed with the Leica Application Suite X (Leica Microsystems Inc. Buffalo Grove, IL). All images were processed using ImageJ 1.44p software^39^ (NIH, Bethesda, MD).

### Western blot

Cells treated with Brazilin and the derivatives were solubilized in ice-cold HEPES-TritonX100 lysis buffer supplemented with Pierce protease and phosphatase inhibitor cocktail (A32961, ThermoScientific^TM^). Cell lysates were quantified with the Bradford (5000205, Bio-Rad) method, and 25 ng of total protein was resolved by 8, 10 or 12% SDS-PAGE and transferred to PVDF membranes (Millipore, Sigma). The membranes were incubated in a Blocking Buffer (37515, ThermoScientific^TM^) for 2 h at RT to block non-specific binding. Membranes were then incubated overnight at 4°C with the primary antibodies for cleaved-caspase 3, cleaved-PARP, Bax, Bcl-2 (1:2000), and GAPDH (1:5000 dilution). The next day, the membranes were washed with TBS/0.1% Tween20 and incubated with a secondary HRP-conjugated anti-rabbit antibody (1:5000 dilution) for 2 h at RT. Proteins were detected using Tanon High-sig ECL Western Blotting substrate (180-5001, ABclonal) in an automated photo-documenter UVP ChemStudio (Analytic-Jena). Images were processed using ImageJ 1.44p software^39^ (NIH, Bethesda, MD).

### Antioxidant activity assays

The antioxidant property of compounds was evaluated by the 2,2-diphenyl-1-picrylhydrazyl (DPPH) radical scavenger activity assay. Briefly, 50 µL of Brazilin or derivatives (2.5-80 µM) were allowed to react with 150 µL of 0.3 mM DPPH-ethanol fresh solution for 30 min at RT protected from light. Ultrapure water and ethanol were used for the blank control. The absorbance was measured at 517 nm in a Multiskan^TM^ GO microplate spectrophotometer (Thermo Fisher Scientific). The percentage of DPPH radical scavenger activity was calculated as [absorbance of control - absorbance of samples)/(absorbance of control)] x100. Results are expressed as half maximal effective concentration (EC_50_); values refer to the concentration (µM) required for 50% of the antioxidant activity.

### Cytoplasmic and mitochondrial oxidative stress evaluation

Dihydroethidium (DHE) and MitoSOX^TM^ assays (Sigma-Aldrich) were performed to evaluate cytosolic and mitochondrial superoxide production, respectively. Briefly, 1x10^4^ cells/cm^2^ were seeded in a 24-well plate and then treated for 48 h with 20 µM and 40 µM of Brazilin and derivatives. After treatments, cells were incubated and protected from light at 37°C with 10 µM DHE or 5 µM MitoSOX^TM^ for 30 and 15 min, respectively. Then, the cells were trypsinized, and the fluorescence was measured using the Spectrum Cellometer (Nexcelom Biosciences) at 370/420 nm (DHE) or 510/580 nm (MitoSOX^TM^) of excitation/emission, as previously described^40^. Data was analyzed with FCS Express 7 (De Novo Software).

### Mitochondrial membrane potential

Changes in mitochondrial membrane potential were determined by the tetramethylrhodamine ethyl ester (TMRE)-mitochondrial membrane potential assay (Abcam, Cambridge, MA). Briefly, 1x10^4^ cells/cm^2^ were seeded and grown in a 24-well plate and then treated for 48 h with 20 µM and 40 µM of Brazilin. After treatments, cells were incubated with 200 nM TMRE for 15 min at 37°C protected from light. FCCP served as a depolarization control and was used as a control for low mitochondrial membrane potential. Cells were trypsinized, and the fluorescence was measured using the Spectrum Cellometer (Nexcelom Biosciences) at 502/595 of excitation/emission, as previously described^40, 41^. Data were analyzed with FCS Express 7 (De Novo Software).

### RNA sequencing and data analysis

Samples for sequencing were obtained from MDA-MB-231 cells treated for 48 h with 20 µM of unmodified Brazilin or derivatives; the RNA was isolated using TRIzol (Invitrogen-Thermo Fisher Scientific), following the manufacturer protocol. Samples were purified by chloroform extraction and isopropanol precipitation, aliquoted for sequencing and RT-qPCR analyses, and flash-frozen in liquid nitrogen and stored at -80 °C until use. Two independent biological replicates were submitted for quality-control analysis, library construction, and sequencing to BGI Genomics **(Supp. Table 3, 4)**. Libraries were sequenced using the BGISEQ-500 platform, and reads were filtered to remove adaptor-polluted, low-quality, and high content of unknown base (N) reads. About 98% of the raw reads were identified as clean reads (∼65 M). The resulting reads were mapped onto the reference human genome (hg38) using HISAT^42^. Transcripts were reconstructed using StringTie^43^, and novel transcripts were identified using Cufflinks^44^ and combined and mapped to the hg38 reference transcriptome using Bowtie2Langmead^45, 46^. Gene expression levels were calculated using RSEM^47^. DEseq2^48^ and PoissonDis^49^ algorithms were used to identify differentially expressed genes (DEG). Gene Ontology (GO) analysis using DAVID (https://david.ncifcrf.gov/tools.jsp) was performed on DEGs to cluster genes into function-based categories.

### RT-qPCR gene expression analysis

Total RNA was purified from three independent biological experiments of MDA-MB-231, MCF7, and MCF10A cells treated for 48 h with 20 µM of unmodified Brazilin or the derivatives. RNA was isolated using TRIzol reagent (Invitrogen-Thermo Fisher Scientific) following the manufacturer’s instructions. cDNA synthesis was performed using 1 µg of RNA as a template and the HiFi Script RT MasterMix (CWBio, Beijing, CN). Quantitative RT-PCR was performed with fast SYBR green master mix (CWBio) on the AriaMX (Agilent Technologies) using the primers listed in **Supp. Table S5**, the delta threshold cycle value (ΔC_T_)^50^ was calculated for each gene and showed as the difference between the C_T_ value of the gene of interest and the C_T_ value of the GAPDH reference gene.

### Statistical analysis

Statistics were performed using ANOVA multiple comparisons with Bonferroni post hoc tests using GraphPad Prism 9.0.2 (La Jolla, CA). A statistical probability of *p<0.05, **p<0.001, ***p<0.001 was considered significant. Independent biological experiments were performed for all analyses.

## ASSOCIATED CONTENT

## Author Contributions

“Conceptualization, M.Z.-E., A.H.-M., M.O., T.P.-B., and N.N.-T.; methodology, M.Z.-E., M.Q., A.H.-M., T. H.-M. and T.S.; validation, M.Z.-E., T. S., M.O., T.P.-B., and N.N.-T.; formal analysis, M.Z.-E., T.S., T.P.-B., and N.N.-T.; investigation, M.Z.-E., M.Q., A.H.-M., T.S., M.O., T.P.-B., and N.N.-T.; resources, M.Z.-E., A.H.-M., T.H.-M., M.O., T.P.-B., and N.N.-T.; data curation, M.Z.-E., T.S., T.P.-B., and N.N.-T.; writing-original draft preparation, M.Z.-E., T.P.-B., and N.N.-T.; review and editing, M.Z.-E., M.Q., A.H.-M., T.S., T. H.-M., M.O., T.P.-B., and N.N.-T.; supervision, M.O., T.P.-B., and N.N.-T.; funding acquisition, M.Z.-E., A.H.-M., T.P.-B., and N.N.-T. All authors have read and agreed to the published version of the manuscript.”

## Data Availability Statement

The RNA-seq datasets are available at GEO. The accession number is: GSE237434.

## Supporting information

Supplemental table 2 DEG

## Acknowledgments

We dedicate this work to the memory of Dr. Napoleon Navarro-Tito, co-senior author of this study, whose insight, mentorship, and vision were instrumental to its conception and execution. We are thankful to Dr. Jorge Bello-Martínez (Universidad Autónoma de Guerrero) for providing us with the initial aliquot of Brazilin isolated and purified from *H. brasiletto*. We also thank Dr. Eugenia Flores Alfaro, Dr. Berenice Illades Aguiar and Dr. Monserrat Olea-Flores for the critical discussions of the work.

