## Supplemental materials for "Cell Death Induced by Homoisoflavonoid Brazilin and its Semi-synthetic Derivates on MDA-MB-231 and MCF7 Breast Cancer Cell Lines"

[†] Deceased. In memory of Dr. Napoleon Navarro-Tito, whose contributions to this work and to the scientific community are deeply appreciated.

[#]Current affiliation: Department of Molecular Biology and Genetics, Johns Hopkins University, School of Medicine, Baltimore, MD 21205

**C.C.S**.

### SUPPLEMENTAL MATERIALS

### Supplementary Figure 1

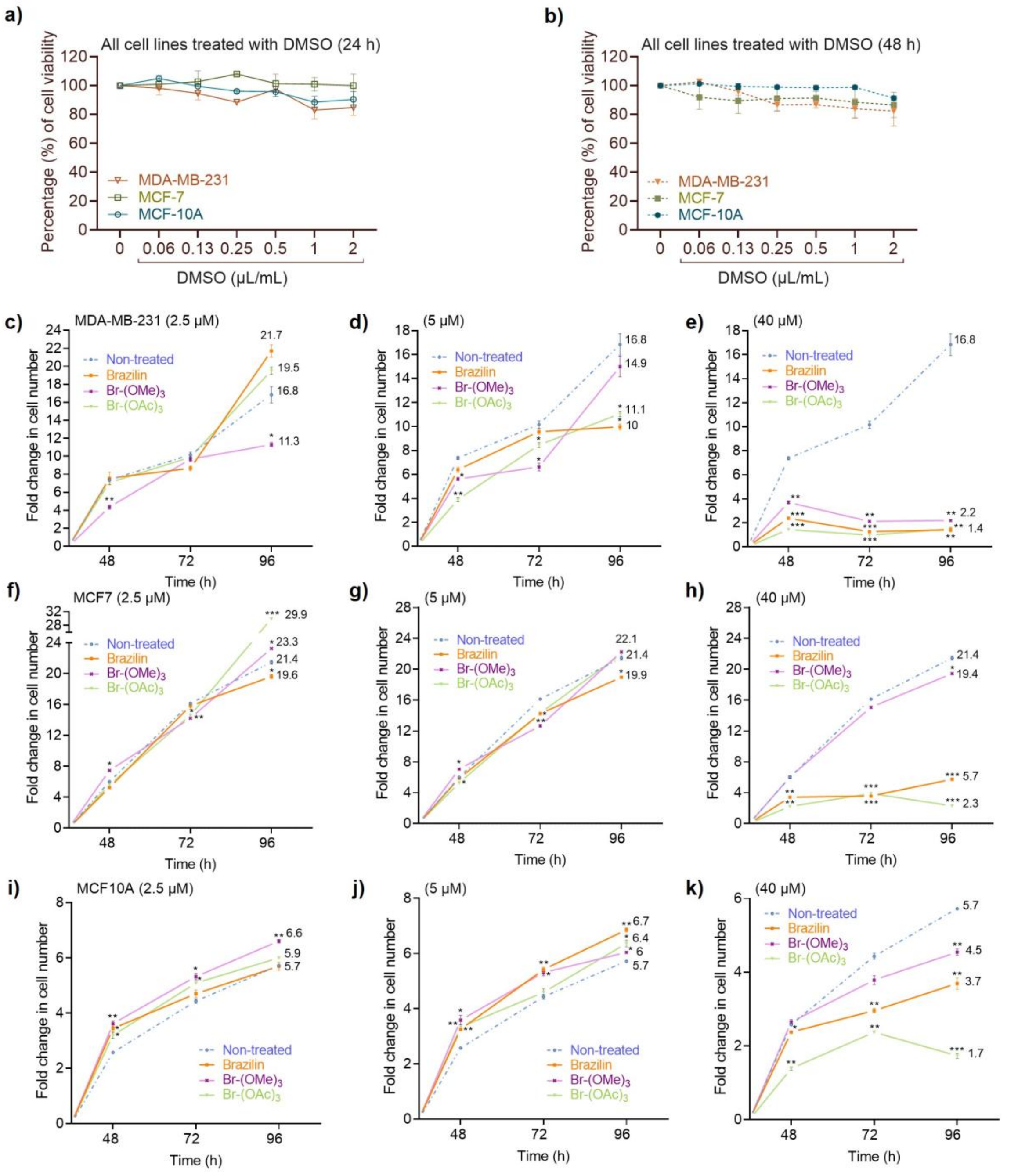


**Supplementary Figure 1. Dose-response effects of DMSO and submaximal concentrations of Brazilin and derivatives on breast cancer and non-tumorigenic cell proliferation.** Serial dilutions of DMSO (µL/mL) were administered to all cell lines, and cell viability was assessed at **(a)** 24 h (solid line) and **(b)** 48 h (dashed line). Cell counting of proliferating cells treated with 2.5 µM, 5 µM, and 40 µM of Brazilin or derivatives is shown for **(c-e)** MDA-MB-231, **(f-h)** MCF7, and **(i-k)** MCF10A cells. Live cells were quantified by trypan blue exclusion at 48 h, 72 h, and 96 h. Growth ratio represents the fold change in live cell number over time and is plotted as mean ± SE. At 96 h, mean values for each treatment are shown. *$p < 0.05$, **$p < 0.001$, ***$p < 0.001$. Data represent three independent biological replicates.

**Supplementary Figure 2**

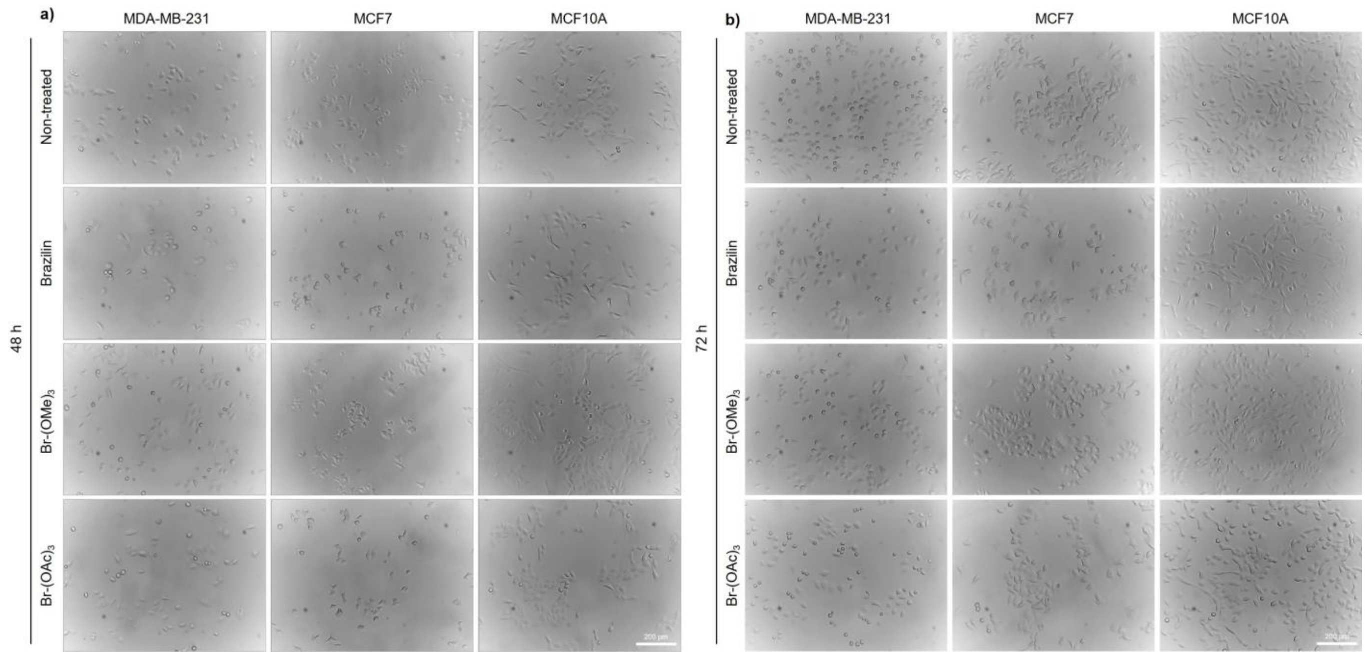


**Supplementary Figure 2. Brightfield imaging of breast cancer and non-tumorigenic epithelial cells during proliferation under Brazilin and derivative treatment.** Representative brightfield images of proliferating **(a)** MDA-MB-231, MCF7, and MCF10A cells after 48 h and **(b)** 72 h of treatment with 20 μM Brazilin or derivatives. Images were captured using a NIKON® ECLIPSE Ts2 microscope. Scale bar = 200 μm. Representative images are shown from three independent biological replicates.

### Supplementary Figure 3

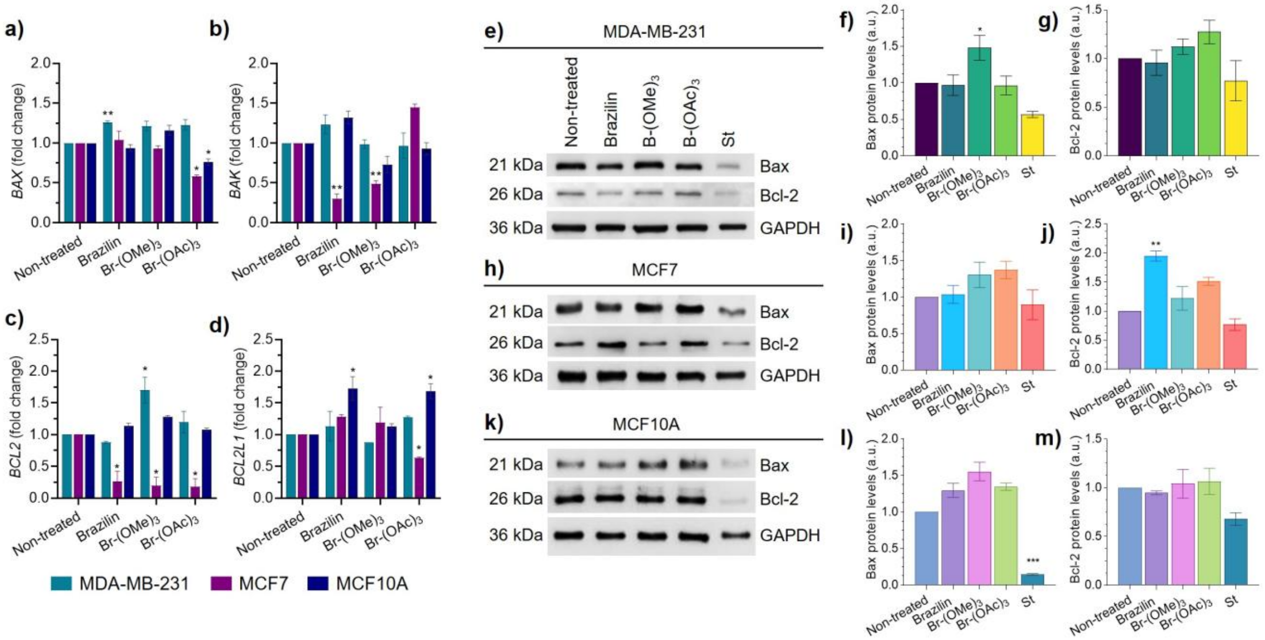


**Supplementary Figure 3. Expression of apoptosis-related genes and proteins in breast cancer and non-tumorigenic cells treated with Brazilin and derivatives. (a-d)** RT-qPCR analysis of BAX, BAK, BCL2, and BCL2L1 mRNA levels in MDA-MB-231, MCF7, and MCF10A cells treated for 48 h with 20 µM Brazilin or derivatives. Parallel mRNA samples used for RNA-seq were used for RT-qPCR. **(e-m)** Immunoblot and quantification of Bax and Bcl-2 protein levels in MDA-MB-231 **(e-g)**, MCF7 **(h-j)**, and MCF10A **(k-m)** cells after 48 h treatment with 20 µM Brazilin or derivatives. 50 nM staurosporine (St) served as apoptosis-positive control. GAPDH was used as loading control. Data are presented as mean ± SE from three independent biological replicates. *p<0.05, **p<0.001, ***p<0.001.

### Supplementary Figure 4

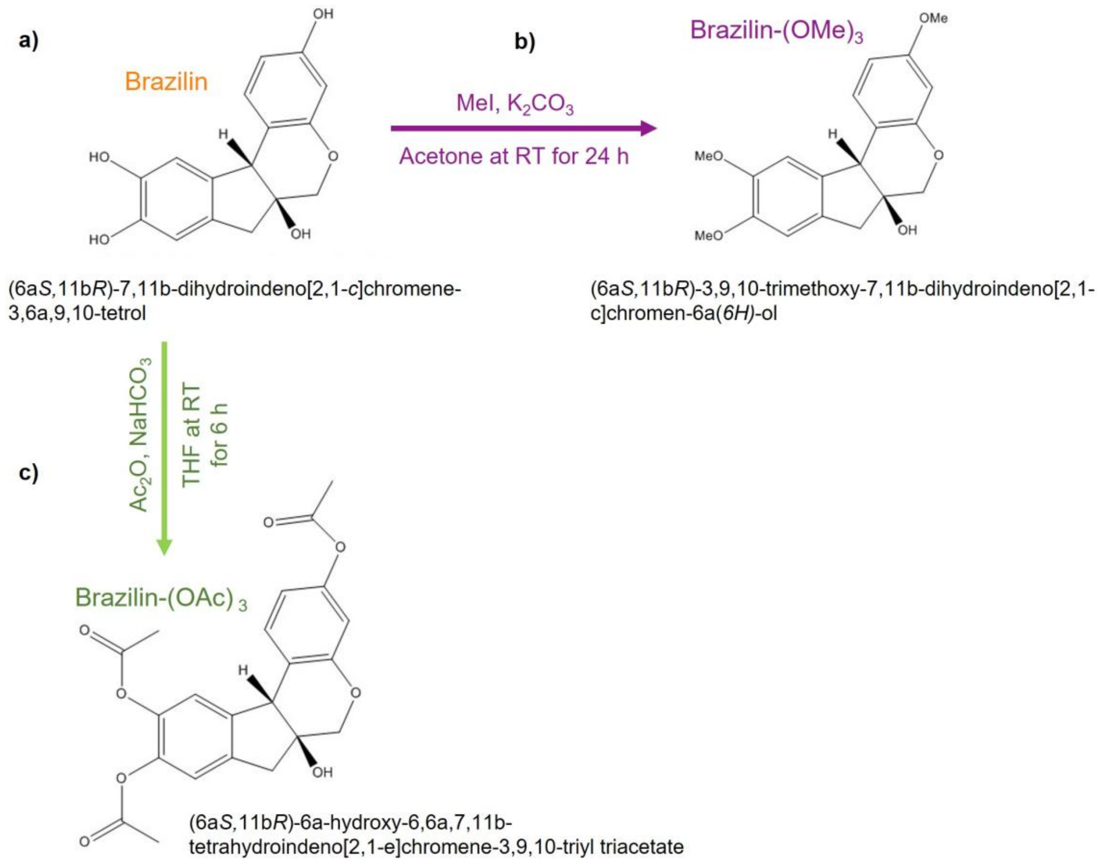


**Supplementary Figure 4. Chemical structures and semi-synthetic routes of Brazilin and derivatives. (a)** Chemical structures of Brazilin, Brazilin-(OMe)$_3$, and Brazilin-(OAc)$_3$. **(b)** Semi-synthesis of Brazilin-(OMe)$_3$: Brazilin (0.14 mmol) in acetone (3.5 mL) was reacted with K2CO3 (0.4191 mmol) and MeI (0.6986 mmol) at room temperature (RT) for 24 h (purple arrow). **(c)** Semi-synthesis of Brazilin-(OAc)$_3$: Brazilin (0.17 mmol) in THF (3 mL) was reacted with $NaHCO_3$ (0.5238 mmol) and $Ac_2O$ (0.8733 mmol) at RT for 6 h (green arrow). Products were purified by flash chromatography (EtOAc:hexane, 65:50) and confirmed by $^1H$ and $^{13}C$ NMR spectroscopy (Hernández-Moreno et al., 2025).

**Supplementary Table 1.** DEG in MDA-MB-231 cells treated with Brazilin and derivatives

| | Total DEG | Upregulated | Downregulated |
|---|---|---|---|
| Non-treated vs Brazilin | 2885 | 1374 | 1511 |
| Non-treated vs Brazilin-$(OMe)_3$ | 11 | 3 | 8 |
| Non-treated vs Brazilin-$(OAc)_3$ | 193 | 120 | 73 |
| Brazilin vs Brazilin-$(OMe)_3$ | 2090 | 1145 | 945 |
| Brazilin vs Brazilin-$(OAc)_3$ | 2611 | 1326 | 1285 |
| Brazilin-$(OMe)_3$ vs Brazilin-$(OAc)_3$ | 8 | 5 | 3 |

**Supplementary Table 3. RNAseq quality statistics**

Data for total reads after filtering and of the clean reads alignments to the reference genes using Bowtie2

| Sample abbreviated name | Sample type | Total raw reads | Total clean reads | Total Mapping Gene Ratio | Uniquely Mapping Gene Ratio |
|---|---|---|---|---|---|
| Control_TNC_1 | Control non-treated cells. Replicate 1 | 94.55 | 92.29 | 84.24 | 80.21 |
| Control_TNC_2 | Control non-treated cells. Replicate 2 | 94.55 | 92.34 | 85.12 | 81.17 |
| TxA_TNC_1 | Brazilin-$(OAc)_3$ TNBC MDA-MB-231 cells Replicate 1 | 92.06 | 89.51 | 83.88 | 79.83 |
| TxA_TNC_2 | Brazilin-$(OAc)_3$ TNBC MDA-MB-231 cells Replicate 2 | 92.06 | 89.61 | 89.61 | 84.70 |
| TxB_TNC_1 | Unmodified-Brazilin TNBC MDA-MB-231 cells Replicate 1 | 89.57 | 87.83 | 84.63 | 80.56 |
| TxB_TNC_2 | Unmodified-Brazilin TNBC MDA-MB-231 cells Replicate 2 | 94.55 | 92.35 | 92.35 | 84.27 |
| TxM_TNC_1 | Brazilin-$(OMe)_3$ TNBC MDA-MB-231 cells Replicate 1 | 94.55 | 92.20 | 85.37 | 81.17 |
| TxM_TNC_2 | Brazilin-$(OMe)_3$ TNBC MDA-MB-231 cells Replicate 2 | 92.06 | 90.03 | 90.03 | 84.22 |

**Supplementary Table 4.** Pearson coefficients RNA-seq

| | Control TNC_1 | Control_ TNC_2 | TxA_T NC_1 | TxA_T NC_2 | TxB_T NC_1 | TxB_T NC_2 | TxM_T NC_1 | TxM_T NC_2 |
|---|---|---|---|---|---|---|---|---|
| Control_ TNC_1 | 1 | 0.997 | 0.967 | 0.965 | 0.969 | 0.971 | 0.97 | 0.99 |
| Control_ TNC_2 | 0.997 | 1 | 0.967 | 0.966 | 0.973 | 0.973 | 0.97 | 0.993 |
| TxA_TN C_1 | 0.967 | 0.967 | 1 | 0.998 | 0.98 | 0.982 | 0.976 | 0.978 |
| TxA_TN C_2 | 0.965 | 0.966 | 0.998 | 1 | 0.982 | 0.982 | 0.98 | 0.977 |
| TxB_TN C_1 | 0.969 | 0.973 | 0.98 | 0.982 | 1 | 0.997 | 0.975 | 0.982 |
| TxB_TN C_2 | 0.971 | 0.93 | 0.982 | 0.982 | 0.997 | 1 | 0.97 | 0.982 |
| TxM_TN C_1 | 0.97 | 0.97 | 0.976 | 0.98 | 0.975 | 0.97 | 1 | 0.97 |
| TxM_TN C_2 | 0.99 | 0.993 | 0.978 | 0.977 | 0.982 | 0.982 | 0.97 | 1 |

**Supplementary Table 5.** Primer sets for RT-qPCR

| Gene | Forward 5'-3' | Reverse 5'-3' |
|---|---|---|
| *CASP3* | GGAAGCGAATCAATGGACTCTGG | GCATCGACATCTGTACCAGACC |
| *BAX* | TCAGGATGCGTCCACCAAGAAG | TGTGTCCACGGCGGCAATCATC |
| *BAK* | TTACCGCCATCAGCAGGAACAG | GGAACTCTGAGTCATAGCGTCG |
| *BCL2* | ATCGCCCTGTGGATGACTGAGT | GCCAGGAGAAATCAAACAGAGGC |
| *BCL2L1* | GCCACTTACCTGAATGACCACC | AACCAGCGGTTGAAGCGTTCCT |
| *SOX4* | GACATGCACAACGCCGAGATCT | GTAGTCAGCCATGTGCTTGAGG |
| *ATF3* | CGCTGGAATCAGTCACTGTCAG | CTTGTTTCGGCACTTTGCAGCTG |
| *GCLM* | TCTTGCCTCCTGCTGTGTGATG | TTGGAAACTTGCTTCAGAAAGCAG |
| *MTRNR2L1* | ACCCAACACAGGCATGCTTA | CTGCACCATTGGGATGTCCT |
| *MTRNR2L2* | GAACCCTCCAAATCCCCCTG | CCGCGGCCGTTAAACATATG |
| *MTRNR2L8* | GCACACCCGTCTATGTAGCA | TTTTGGTAAACAGGCGGGGT |
| *GAPDH* | GCCATCAAGGAGGCTGTAAAAGC | GGTATCACCGAGGAAGTCCGTA |
